# A Red Fluorescent Lifeact Marker to Study Actin Morphology in Podocytes

**DOI:** 10.1101/2024.09.17.613449

**Authors:** Eva Wiesner, Julia Binz-Lotter, Simon E. Tröder, David Unnersjö-Jess, Nelli Rutkowski, Branko Zevnik, Thomas Benzing, Roland Wedlich-Soldner, Matthias J. Hackl

## Abstract

F-actin is a major component of the cellular cytoskeleton, responsible for maintaining cell shape, enabling movement and facilitating intracellular transport. In the kidney, glomerular podocytes are highly dependent on their actin cytoskeleton shaping their unique foot processes. Hereditary mutations in actin-binding proteins cause focal segmental glomerulosclerosis, while other organs remain largely unaffected.

So far, actin visualization in podocytes has been limited to electron microscopy or indirect immunofluorescent labeling of actin-binding proteins. However, the short F-actin-binding peptide Lifeact enables researchers to study actin dynamics *in vitro* and *in vivo* with minimal interference with actin metabolism.

Here we introduce a new mouse model with conditional expression of a Lifeact.mScarlet-I fusion protein providing red labeling of actin. Cre recombinase-mediated activity allows cell-specific and mosaic expression in podocytes, enabling selective labeling of individual cells to contrast with non-expressing neighboring cells. Transgenic mice are born healthy and young animals display no kidney-related phenotype. By intravital imaging and super-resolution microscopy, we show subcellular localization of actin to the foot processes in a resolution previously only obtainable by electron microscopy.

Our novel mouse line provides the opportunity to study the actin cytoskeleton in podocytes and other cell types by intravital imaging and other conventional light microscopy techniques.

## Introduction

The glomerulus forms the functional filtration unit of the kidney. Glomeruli filter plasma through a complex filtration barrier resulting in a protein-free primary urine. The filter consists of a fenestrated endothelial cell layer, the glomerular basement membrane, and a layer of epithelial cells called podocytes. Podocytes have a large cell body with protrusions termed primary processes, which further divide into smaller foot processes (FPs). FPs of neighboring podocytes interdigitate and are connected by a specialized cell-cell contact, the slit diaphragm (SD). In the SD, nephrin is one of the most prominent proteins. A sophisticated network of tubulin in the primary processes and actin bundles in the FPs support the complex podocyte structure.[1]

During disease, the actin cytoskeleton drastically rearranges, leading to flattening and simplification of FPs structure, a process termed ‘foot process effacement’.[2] Human congenital mutations in genes encoding actin-interacting proteins such as alpha-actinin-4, podocin and CD2AP result in kidney disease, while other organs show less or no phenotype, and thus highlight the significance of an intact cytoskeleton for podocyte function.[3],[4],[5] Therefore, visualization of podocyte actin cytoskeleton has been increasingly important to understand disease development and evaluate podocyte injury.[2]

In 2008 Riedl et. al. published Lifeact as a novel actin-binding probe. Lifeact is a 17 amino acid long peptide and binds preferentially to filamentous actin (F-actin).[6] Since its publication, Lifeact has been fused to a variety of fluorescent proteins to label actin *in vitro* and *in vivo*. Tissue inducible expression and constitutive expression of Lifeact.EGFP and Lifeact.mRuby in mice resulted in viable, fertile and phenotypically normal animals.[7],[8]

Transient expression of Lifeact in cell culture did not alter actin dynamics.[9] Detailed structural analysis of Lifeact binding to actin using cryoEM revealed no interference with the helical symmetry of the filament.[10] Super-resolution imaging of Lifeact and phalloidin in HeLa cells showed comparable results for imaging quality, filament thickness and length.[11] However, Lifeact does not bind all actin forms equally, but shows a higher binding affinity towards F-actin.[12]

To date, only a conditional Lifeact.EGFP mouse line has been generated.[8] This limits its combination with any other green reporter proteins such as GCaMP, due to overlapping emission spectra. We therefore generated a novel conditional Lifeact mouse line expressing tissue-specific Lifeact tagged with the red fluorescent protein mScarlet-I. In the group of monomeric red fluorescent proteins, mScarlet-I displays the highest calculated brightness at its fluorescence emission maximum at 569 nm, with a more than 3.5-fold increase compared to mCherry.[13] In recent years, the development of super-resolution techniques such as STED and SIM have enabled visualization of podocyte architecture similar to electron microscopy.[14][15] Here, we demonstrate efficient endogenous actin labeling in podocytes in transgenic mice. Immunofluorescent co-staining with other proteins provides further insight into cytoskeletal structures.

## Material & Methods

### Generation of targeting vector

For construction of the plasmid pR26 CAG LifeAct.mScarlet-I the sequence of LifeAct.mScarlet-I was cloned from the plasmid pLifeAct_mScarlet-i_N1 (Addgene #85056) in the backbone of the plasmid pR26 CAG AsiSI/MluI (Addgene #74286) using the restriction enzymes AsiSI and MluI.[13] [16] DNA Sequence of Lifeact.mScarlet-I was amplified using the forward primer 5-<u>GCGATCGC</u>ATTCCACCATGGGCGTGGCCG-3’ including an <u>AsiSI</u> restriction site and a reverse primer 5-CGCG<u>ACGCGT</u>CTTGTACAGCTCGTCCATGCCGCC-3’ including a <u>MluI</u> restriction site. After purification, the PCR fragment was digested and ligated into the equally digested pR26 CAG AsiSI/MluI plasmid to obtain the construct pR26 CAG LifeAct.mScarlet-I.[16]

### Mouse genome editing

LifeAct.mScarlet-I (C57BL/6N-Gt(ROSA)26Sor^em333(CAG-mScarlet-I)Cecad^/Cecad) mice for Cre-recombinase mediated expression of LifeAct.mScarlet-I from the Gt(ROSA)26Sor locus were generated by CRISPR/Cas9 genome editing using a well-established gene targeting strategy.[16] Therefore, 400 nM of each Alt-R™ guide RNA and 20 ng/µl plasmid DNA repair template (pR26 CAG LifeAct.mScarlet-I) were injected with 200 nM Alt-R™ SpCas9 protein and 30 ng/µl SpCas9 mRNA (TriLink, L-6125-20) in C57BL/6NRj zygotes as previously published.[17] CRISPR reagents were purchased from Integrated DNA Technologies (USA) and animals were obtained from Janvier Labs (France). Embryos were cultured *in vitro* and transferred to recipients as described.[18] Genome editing was performed at the *in vivo* Research Facility of the CECAD Research Center, University of Cologne, Germany.

### Genetic validation of Knock-In

For a PCR to validate the integration of our construct into the Rosa26 locus as described by Chu et. al., we used the primers SAR CCTGGACTACTGCGCCCTACAGA und R26F3 CTGCCCGAGCGGAAACGCCACTGAC.[16] The PCR was carried out using the Herculase II Fusion DNA Polymerase (Agilent Technologies, #600675) resulting in a ∼ 1.3 kb fragment. After successful integration validation, the genetic locus was further analyzed by Cergentis. Following their protocol (www.cergentis.com) bone marrow of heterozygous mice was prepared and analyzed. Two sets of primers (see table 1) were used for targeted Locus Amplification (TLA) sequencing.

**Table 1.**
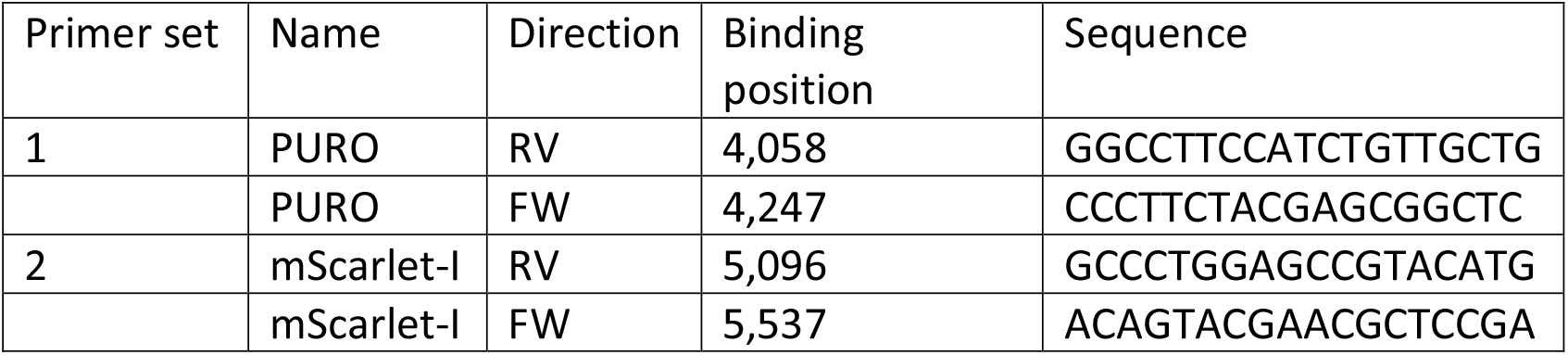
TLA Sequencing Primers.

### Mice

Animals were derived and maintained in the CECAD Research Center, University of Cologne, Germany, in individually ventilated cages (Greenline GM500; Tecniplast) at 22°C (± 2°C) and a relative humidity of 55% (± 5%) under 12-hour light cycle on sterilized bedding (Aspen wood, Abedd, Germany) and with access to sterilized commercial pelleted diet (Ssniff Spezialdiäten GmbH) and acidified water *ad libitum*. The microbiological status was examined as recommended by the <u>F</u>ederation of <u>E</u>uropean <u>L</u>aboratory <u>A</u>nimal <u>S</u>cience <u>A</u>ssociations (FELASA) and the mice were free of all listed agents including opportunists. Mice (C57BL/6N) of both sexes at the age of 4 weeks and 62-weeks were used. Generated C57BL/6N-Gt(ROSA)26Sor^em333(CAG-mScarlet-I)Cecad^/Cecad (short: Lifeact.mScarlet-I)) were crossed to podocin-Cre (MGI: 3586451) to achieve expression in all podocytes (Pod^Lifeact.mScarlet-I^). Inducible expression was obtained by using the tamoxifen responsive podocin-iCreERT2 (MGI: 4819592) to generate iPod^Lifeact.mScarlet-I^.[19], [20] Cre negative animals (Pod^wt^) were used as control. For phenotypic analysis, 3 mice per age with the genotype Pod^Lifeact.mScarlet-I^ were used. As control 2 Pod^wt^ per age group were used. No animals were excluded, no randomization was performed. Potential confounders were not controlled and blinding of the study was not performed. N equals the number of mice if not stated otherwise. For measurement of SD length/area, 5 glomeruli per animal were imaged and analyzed.

### Genotyping protocol

For genotyping, we used skin biopsies obtained by mouse ear tagging. DNA was extracted using the HotSHOT protocol and amplified using REDTaq ReadyMix (Sigma-Aldrich).[21] For amplification of the Rosa26 wildtype allele using the forward primer GGAGTGTTGCAATACCTTTCTGGGAGTTC and reverse primer CGAGGCGGATCACAAGCAATA results in a ∼ 350 bp band. The Rosa26 transgenic locus was amplified using the forward primer GGGGAGGATTGGGAAGACAA together with CGAGGCGGATCACAAGCAATA as the reverse primer resulting in a 560 bp band. Genotyping for Podocin-Cre and podocin-iCreERT2 was performed as previously published.[19], [20]

### Analysis of proteinuria and creatinine

Proteinuria was evaluated by measuring albumin content in urine samples. In short, 2 µl of urine sample were mixed with 8 µl H_2_O and 2 µl 6x Laemmli-loading dye. Samples were boiled (5 min, 95°C) and loaded onto a 10 % SDS gel. As a reference for protein content 1 µg and 10 µg bovine serum albumin (BSA) were used. After gel electrophoresis, the gel was stained with Coomassie Brilliant Blue to visualize albumin bands. For a urinary albumin ELISA, a 96-well plate was covered with an *α*-mouse albumin antibody (1:10.000) for 1 h at RT (A90-134A, Bethyl Laboratories). The plate was washed 5 times and incubated with a blocking solution for 30 min followed by further 5 washing steps. An albumin standard was prepared for a reference curve ranging from 0 to 625 ng/ml. Urines were diluted to a concentration of around 150 ng/ml. Standard and urine samples were added to the plate and removed after 1 h. For detection, an HRP-coupled goat *α*-mouse antibody (1:25.000) was used (# A90-134P, Bethyl Laboratories). Ultimately, the absorbance at 450 nm was measured on a plate reader (Perkin Elmer).

Urinary creatinine was determined using a commercially available kit (Creatinine (urinary) Colorimetric Assay Kit, Item No. 500701, Cayman Chemical).

### Tamoxifen Induction

LifeAct.mScarlet-I^fl/wt^ ERT2.Pod:cre^tg/wt^ (iPod^LifeAct.mScarlet-I^) mice were induced with Tamoxifen (0.1 mg/g body weight) in sesame oil via oral gavage at the age of 3 weeks. Control animals received sesame oil via oral gavage. Mice were monitored for weight loss and sacrificed 5 days after induction to either prepare acute kidney slices or used for *in vivo* imaging.

### In vivo imaging

Preparation and *in vivo* imaging were performed as previously published.[22] Before surgery mice received buprenorphine (RB Pharmaceuticals Limited, 0.1 mg/g BW) as an analgesic drug and were anesthetized with isoflurane (Piramal Critical Care, #9714675, 1.5-5 %). A 1 cm long incision in the midline of the neck was made and the right carotid artery was dissected. The artery was ligated cranially and clamped caudally. A small incision was made, and a catheter (PE-10) was inserted and fixed with sutures. Next, the left kidney was exteriorized through a small incision above the kidney and immobilized with a custom-build kidney holder (Medres). A coverslip was placed on the kidney with minimum pressure. A 70-kDa fluorescein coupled dextran (D1823, Invitrogen) was administered through the catheter to label the blood circulation. The body temperature was maintained with a heating system (Medres). *In vivo* imaging was carried out using an upright multiphoton microscope (TCS SP8 MP-OPO; Leica Microsystems) and a 25x water immersion objective (numerical aperture: 0.95). The multiphoton laser (Chameleon Vision II; Coherent) was tuned to 940 nm and images were recorded using non-descanned hybrid detectors and a FITC/TRITC (525/50) filter set. Stacks of glomeruli were recorded with a 2 µm step size. Laser power was adjusted to the brightness of the observed podocytes to minimize saturation. At the end of the experiment, mice were sacrificed by cervical dislocation, while still under anesthesia.

### Preparation and imaging of acute kidney slices

Fresh mouse kidneys were prepared as previously described.[23] Shortly, kidneys were embedded in low-melting agarose (4%) and cut into 300 µm slices using a vibratome (VT1200 S, Leica). Slices were maintained in Krebs-Henseleit-Buffer carbonated with 95% O_2_/5% CO_2_.[24] For imaging, single slices were transferred into an imaging dish and stabilized with a slice anchor (Warner Instruments) to reduce tissue drifting. Multiphoton imaging was performed with a TCS SP8 MP-OPO microscope (Leica Microsystems) with a Chameleon Vision II laser (Coherent) at 940 nm.

### STED sample preparation and imaging

Immunofluorescent labeling was conducted as previously described.[14] Briefly, 4 % PFA-fixed kidney tissue was immersed in a hydrogel solution. The gel was polymerized for 3 h at 37°C. Then, the tissue was embedded in 3% agarose and cut into 200 µm thick slices using a vibratome (VT1200 S, Leica). The slices were optically cleared overnight, followed by incubation with target protein antibodies (see Table 2). Confocal and STED images of fixed slices were acquired with a Leica TCS SP8 gSTED 3X equipped with a white light laser (WLL) and a 775 nm STED depletion laser (Leica Microsystems).

**Table 2.** Antibodies used for immunofluorescent stainings.

| **Antibody** | **dilution** | **Cat no. & company** |
| --- | --- | --- |
| Synaptopodin (SE-19) | 1:200 | S9442-200UL, Sigma |
| tdTomato | 1:200 | AB8181-200, Sicgen |
| Human Nephrin Antibody | 1:100 | AF4269, R&D Systems |
| RFP | 1:200 | 600-401-379, Rockland |
| Anti-Tubulin, Acetylated | 1:200 | T6793-100UL, Sigma |
| Vimentin (D21H3) XP<br>(Alexa488 conjugated) | 1:100 | 9854S, Cell Signaling |
| GFP (Alexa647 conjugated) | 1:100 | A-31852, Invitrogen |
| FluoTag®-X2 anti-mScarlet-I<br>(AbberiorStar635P conjugated) | 1:50 | N1302-Ab635P-L, NanoTag<br>Biotechnologies |
| **Secondary antibody** |  |  |
| donkey anti-rabbit Alexa488 | 1:100 | Jackson ImmunoResearch |
| donkey anti-goat<br>AbberiorStar635P | 1:100 | Jackson Immunoresearch,<br>coupled in house |
| donkey anti-goat Atto594 | 1:100 | Jackson Immunoresearch,<br>coupled in house |
| donkey anti-rabbit Atto594 | 1:100 | Jackson Immunoresearch,<br>coupled in house |
| donkey anti-rabbit<br>AbberiorStar635P | 1:100 | Jackson Immunoresearch,<br>coupled in house |
| donkey anti-mouse Alexa488 | 1:100 | Jackson ImmunoResearch |

### Image processing

Images were processed using FIJI and Omero.figure web application. Movie of 3D rendering of z-stack was created using Imaris.

### Statistical analysis

For statistical analysis of data, the software Graph Pad Prism (Version 10.1.2) was used. We performed an ordinary two-way ANOVA with multiple comparisons for all performed tests in Fig.2. No data points were excluded. ns p > 0.05, * p ≤ 0.05, ** p ≤ 0.01, *** p ≤ 0.001, **** p ≤ 0.0001.

## Results

To generate a conditional Lifeact.mScarlet-I mouse line, the desired sequence was cloned into the ROSA targeting vector pR26 CAG AsiSI/MluI.[16] The plasmid was then targeted with CRISPR/Cas9 genome editing to the Gt(ROSA)26Sor locus as previously described (Fig. 1a).[16] Integration PCR performed on the 9 founder animals (#1-9) resulted in one positive result (#7) with a band at 1.3 kB (Fig. 1b). Offspring of this mouse were then used to establish the mouse line. We then performed TLA and sequencing of F_1_ mice to verify the correct sequence and integration of the insert as well as exclude unspecific vector integration. TLA sequencing showed correct integration in the Gt(ROSA)26Sor locus on chromosome 6 between exons 1 and 2 without additional integration sites in the entire genome (Fig. 1c). Apart from the notoriously complex CAG promotor sequence, TLA sequencing also confirmed the sequence of the entire insert without any mutation (Suppl. Fig. S1).

**Figure 1.**
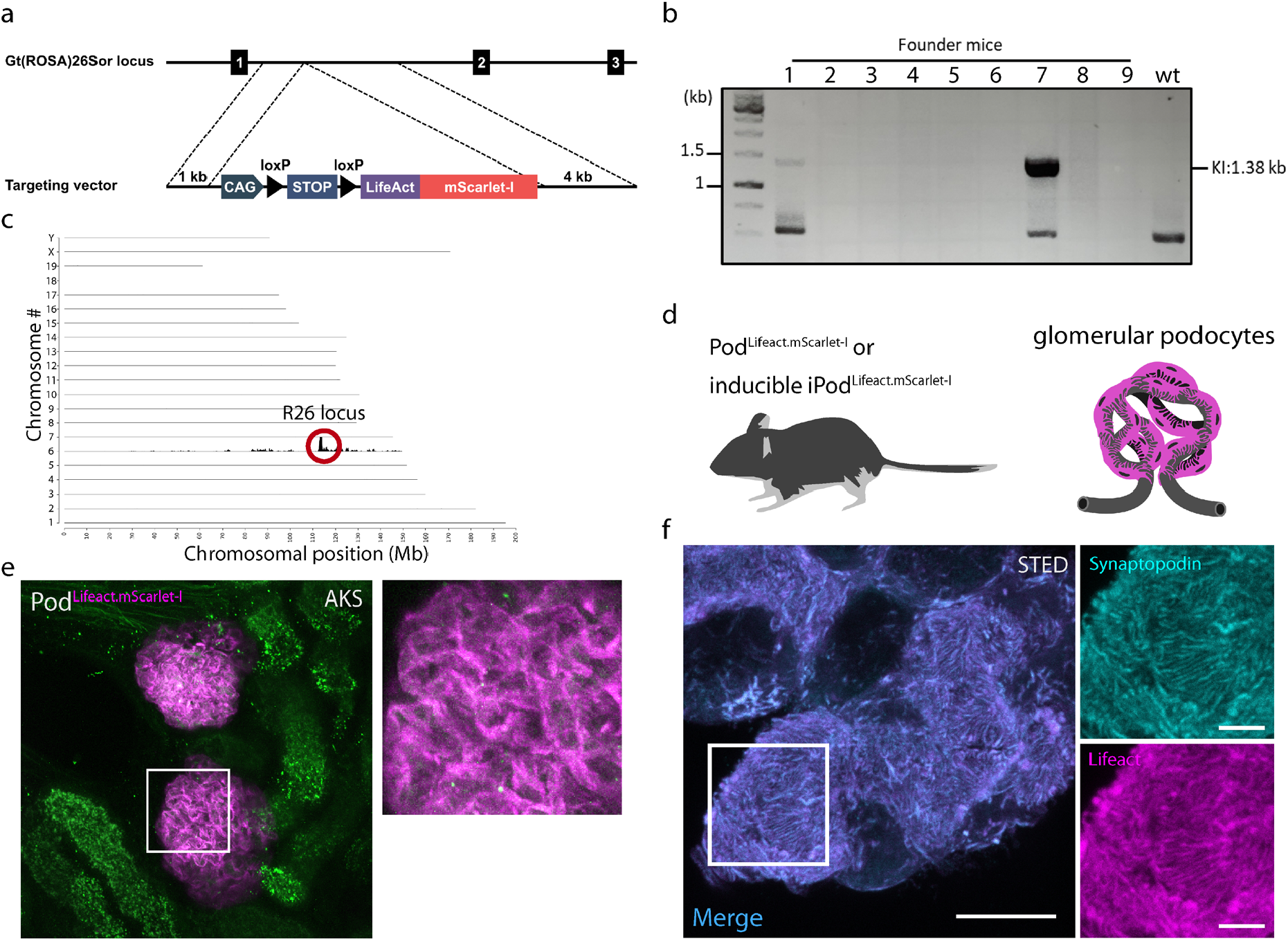
Generation of Lifeact.mScarlet-I mice to label filamentous actin. **(a)** Targeting strategy of the ROSA26 locus for site-specific integration of R26 LSL Lifeact.mScarlet-I to mediate Cre dependent expression. **(b)** Integration PCR of different founder mice to validate successful integration in the F_0_ generation. **(c)** TLA sequence coverage across the mouse genome using primer set 2. The chromosomes are indicated on the y-axis, the chromosomal position on the x-axis. Identified integration site is encircled in red. **(d)** The Lifeact.mScarlet-I construct was expressed in glomerular podocytes using podocin-cre (Pod^Lifeact.mScarlet-I^) or tamoxifen inducible podocin-Cre-ERT2 (iPod^Lifeact.mScarlet-I^). Created with BioRender. **(e)** Maximum intensity projection (MIP) image of Lifeact.mScarlet-I fluorescence in podocytes in murine acute kidney slice (AKS). **(f)** Representative MIP STED microscopy images of Pod^Lifeact.mScarlet-I^ glomerular podocytes co-stained with synaptopodin. Annotation: KI -Knock-In, TLA – targeted locus amplification, MIP – maximum intensity projection, AKS – acute kidney slice. Scale bar – 10 µm, zoom in – 3 µm.

We then used Pod-Cre (Pod^Lifeact.mScarlet-I^) or tamoxifen inducible Pod-ERT2-Cre (iPod^Lifeact.mScarlet-I^) mice to express Lifeact.mScarlet-I specifically in podocytes (Fig. 1d). Lifeact.mScarlet-I expression in heterozygous Pod^Lifeact.mScarlet-I^ animals resulted in bright podocyte specific fluorescence in acute kidney slices (AKS) (Fig. 1e). In fixed tissue Lifeact.mScarlet-I completely co-localized with the actin binding protein synaptopodin (Fig. 1f).

To exclude the occurrence of adverse effects of Lifeact.mScarlet-I expression on kidney function due to binding to the actin cytoskeleton we investigated young mice at the age of 4-weeks and aged mice at 62-weeks of age. We therefore compared heterozygous Pod^Lifeact.mScarlet-I^ animals to Cre negative littermates (Pod^wt^).

Spot urine samples revealed no albuminuria in young mice, while aged Pod^Lifeact.mScarlet-I^ animals showed mild proteinuria in their urine samples compared to same-age Pod^wt^ littermates (Fig. 2a). This is also reflected by the increased albumin creatinine ratio (ACR) in 62-week old Pod^Lifeact.mScarlet-I^ mice, while 4-week old mice of both genotypes and 62-week old Pod^wt^ had low ACRs (Fig. 2b). However, serum creatinine levels in young Pod^wt^ and Pod^Lifeact.mScarlet-I^ mice were higher compared to the aged mice of both genotypes (Suppl. Fig. S2). To evaluate the effects of Lifeact.mScarlet-I expression on SD structure, we performed stimulated emission depletion (STED) microscopy to visualize nephrin expression at the SD (Fig. 2c). Young Pod^Lifeact.mScarlet-I^ mice showed a small reduction of the SD length/area compared to Pod^wt^ animals (Fig. 2d). Old Pod^Lifeact.mScarlet-I^ mice presented with a significant further reduction in SD length/area compared to old Pod^wt^ animals, indicating a mild effect of LifeAct.mScarlet-I expression on SD integrity during aging (Fig. 2d).

**Figure 2.**
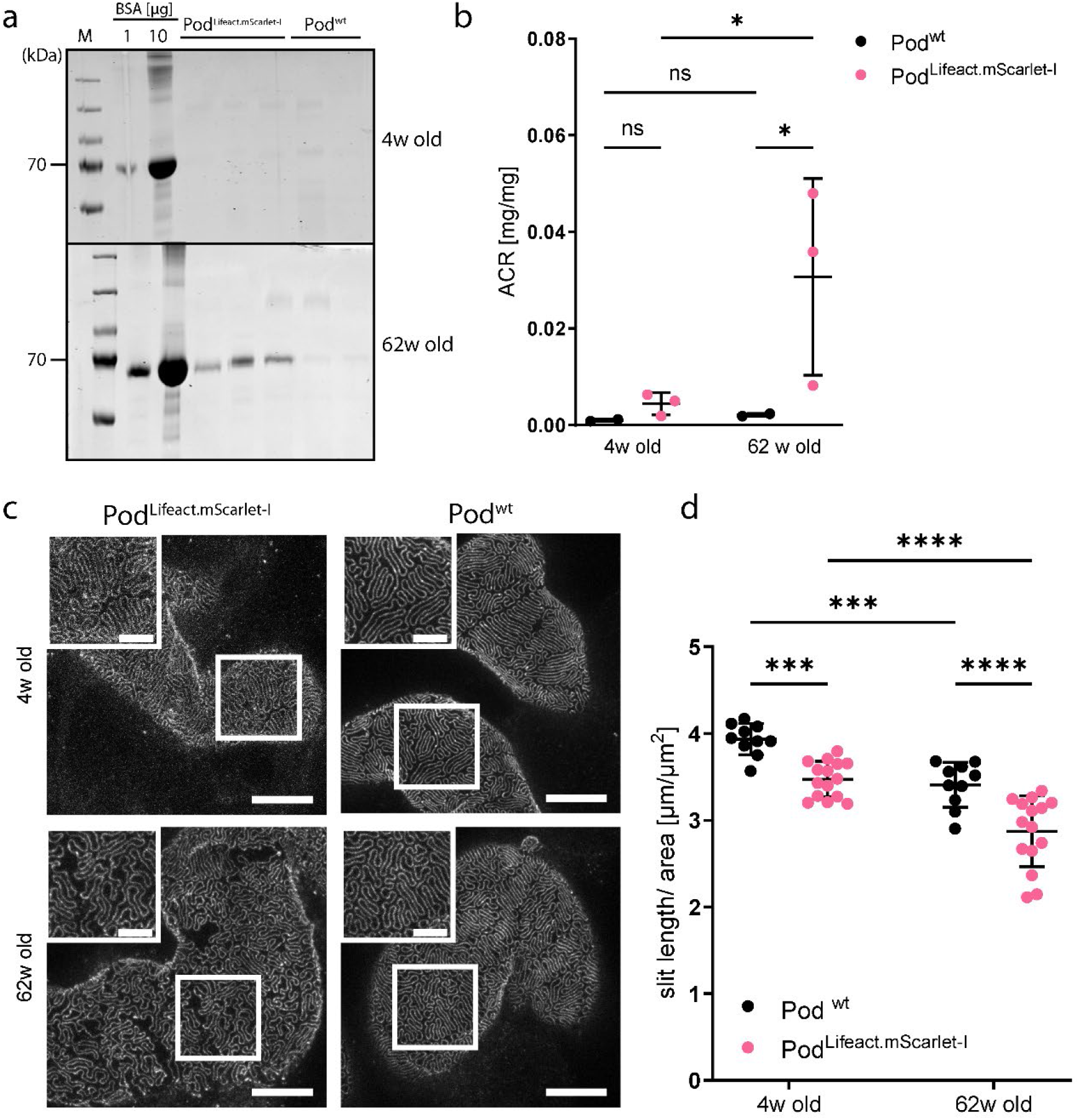
Phenotypic analysis of young and aged Lifeact.mScarlet-I mice. **(a)** Coomassie urine analysis of 4- and 62-week old Pod^Lifeact.mScarlet-I^ (n=3 per age) and Pod^wt^ (n=2 per age) mice. 1 and 10 µg bovine serum albumin (BSA) was used as a standard. **(b)** Urinary albumin/creatinine ratio from spot urine samples of 4-& 62-week old Pod^Lifeact.mScarlet-I^ (4-weeks: n=3; 62 weeks: n=3) and Pod^wt^ (4-weeks: n=2; 62 weeks: n=2) mice. **(c)** Representative images of nephrin expression in 4- and 62-week old Pod^Lifeact.mScarlet-I^ and Pod^wt^ mice using STED microscopy. **(d)** Slit diaphragm length/area measurements of 4- and 62-week old Pod^Lifeact.mScarlet-I^ (n=15 glomeruli of three mice each) and 4- and 62-week old Pod^wt^ (n=10 glomeruli of two mice each). Scale bar – 5 µm, zoom in – 2 µm. Statistical analysis: Ordinary two-way ANOVA with multiple comparisons. ns p > 0.05, * p ≤ 0.05, ** p ≤ 0.01, *** p ≤ 0.001, **** p ≤ 0.0001.

Subsequently, we performed intravital imaging of the kidney to evaluate the Lifeact expression pattern in 4-week old Pod^Lifeact.mScarlet-I^ animals *in vivo*. Glomerular blood flow was labeled by injecting a FITC conjugated 70 kDa dextrane. A video of the acquired z-stack shows capillary blood flow (Suppl. Video V1). Pod^Lifeact.mScarlet-I^ mice showed a homogenous expression of Lifeact.mScarlet-I in all podocytes as depicted in the maximum intensity projection (MIP) (Fig. 3a). Single plane images show strong Lifeact.mScarlet-I fluorescence close to the capillary which indicates labeling of FPs (Fig.3 a’ & a”). Next, we prepared acute kidney slices (AKS) to establish Lifeact.mScarlet-I as an *ex vivo* tool to study the cytoskeleton.[23] Comparable to podocyte *in vivo* imaging, glomeruli of Pod^Lifeact.mScarlet-I^ mice showed homogenous expression of Lifeact.mScarlet-I adjacent to the capillary (Fig. 3b-b”). Finally, we used the Lifeact.mScarlet-I mouse model to visualize the actin cytoskeleton in FPs by super-resolution microscopy. Staining of mScarlet-I using an α-RFP antibody showed a homogenous staining pattern in Pod^Lifeact.mScarlet-I^ mice. The MIP shows intense labeling of the capillaries and minimal labeling of the cytoplasm within the cell body (Fig. 3c-c’). STED images of the capillary loops show a densely packed layer of FPs which covers the entire surface in a regular pattern (Fig. 3c”). The SD is visible as an unstained line between single FPs. 3D rendering of a z-stack revealed an actin distribution with pronounced ridges between single FPs wrapping around the glomerular capillary (Suppl. Video V2).

**Figure 3.**
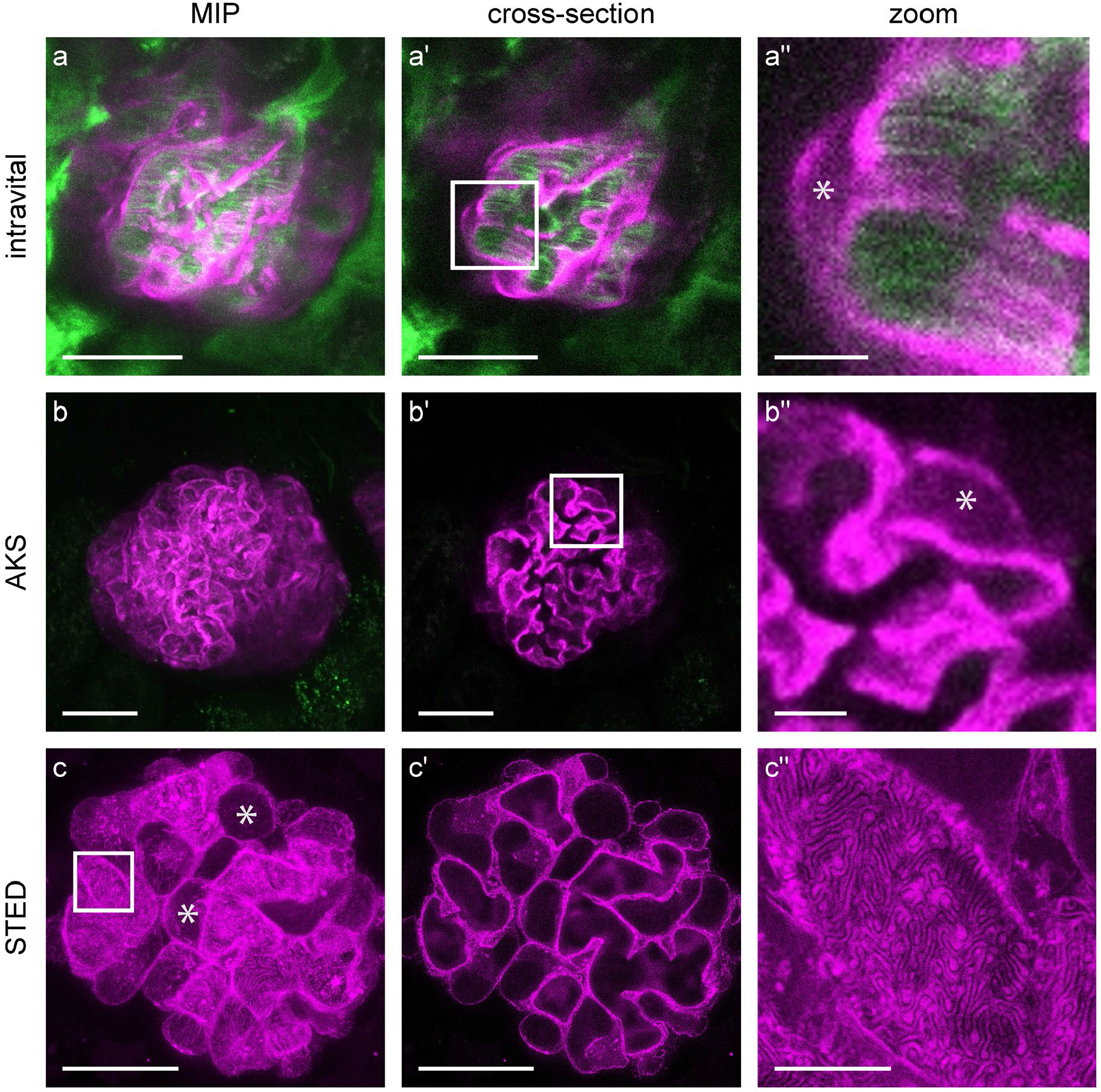
Homogenous expression of Lifeact.mScarlet-I in glomerular podocytes. **(a)** Representative images of Lifeact.mScarlet-I fluorescence in podocytes imaged by intravital microscopy using a multiphoton microscope (magenta). Blood flow is visualized by injection of 75-kDa FITC coupled dextran (green). **(b)** Murine AKS of Pod^Lifeact.mScarlet-I^ animals imaged with multiphoton microscopy. **(c)** Immunofluorescently labeled Lifeact.mScarlet-I tissue section imaged with STED microscopy. Asterisks indicate podocyte cell bodies. Scale bars – 25 µm, zoom – 5 µm; MIP – maximum intensity projection; AKS – acute kidney slice.

We validated the specificity of RFP or tdTomato antibodies for mScarlet-I by co-staining with an anti-mScarlet-I nanobody (Suppl. Fig. S3). Even though the nanobody penetrated deeper into the tissue, high concentrations were needed to obtain a similar staining intensity.

Next, we prepared AKS from Pod^Lifeact.mScarlet-I^ mice to test the possibility of visualizing dynamic actin reorganization. We, therefore, incubated AKS of Pod^Lifeact.mScarlet-I^ with increasing doses of methyl-β-cyclodextrin (MβCD). MβCD depletes cholesterol from plasma membranes and disrupts lipid rafts and thus the SD complex. Figure 4a shows a representative image of a control glomerulus without MβCD with a normal distribution of Lifeact.mScarlet-I. After incubation with 20 mM MβCD for 4 h the regular actin pattern is partially disrupted (Figure 4b) and actin blebbing is visible in AKS (Fig. 4c’, arrow). After fixation, the observed actin blebs remained and a reduction of the continuous coverage of the capillary with actin cables became evident (Fig. 4d’, arrow). Noticeably cells detached from the capillary surface during processing of the sample indicated by asterisks, which was not the case in control animals (Fig. 4d). In areas of podocyte detachment the SD pattern was completely disrupted.

**Figure 4.**
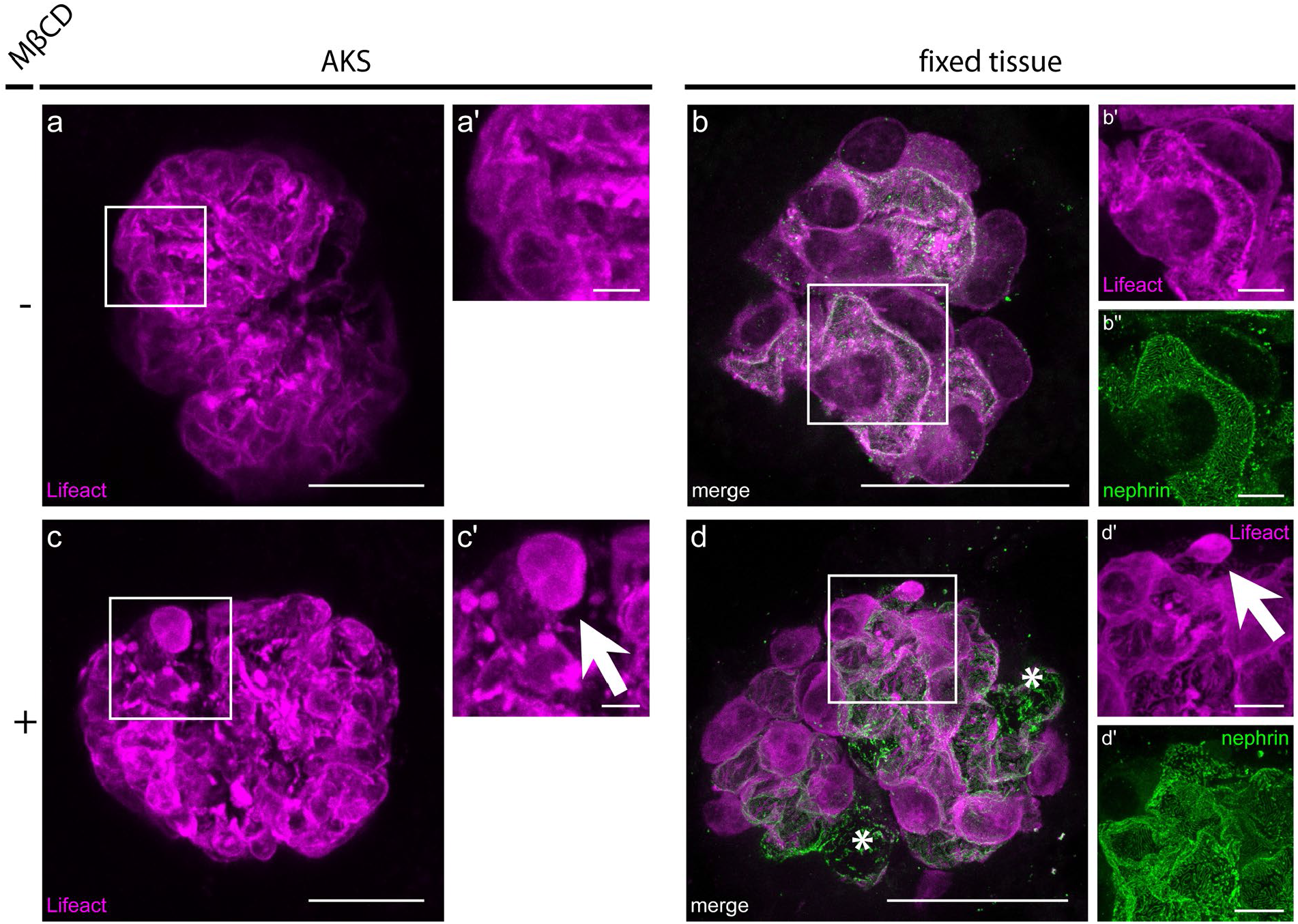
Depletion of lipid rafts leads to actin disruption in Pod^Lifeact.mScarlet-I^acute murine kidney slices. **(a)** Representative image of Lifeact.mScarlet-I pattern (magenta) in control conditions using multiphoton microscopy. **(b)** Representative images of Lifeact.mScarlet-I (magenta) and nephrin pattern (green) in control conditions in fixed tissue using STED microscopy. **(c)** Representative image of a glomerulus after incubation with 20 mM MβCD for 4h imaged with multiphoton microscopy. Arrow in zoom-in indicates actin blebbling (c’). **(d)** Representative image of a Lifeact.mScarlet-I (magenta) and nephrin pattern (green) after incubation with 20mM MβCD for 4h in fixed tissue imaged with STED microscopy. Arrow in zoom-in indicates actin blebbling, asterisk indicates areas with detached podocytes. Scale bars – 25 µm, zoom – 5 µm. MβCD – Methyl-β-cyclodextrine.

To enhance contrast during imaging we used a tamoxifen inducible Pod-ERT2-Cre (iPod^Lifeact.mScarlet-I^) mouse line to achieve Lifeact.mScarlet-I expression only in a subset of podocytes. iPod^Lifeact.mScarlet-I^ mice (0.1 mg/g tamoxifen, 5 days before imaging) revealed partial expression of the reporter. Thus, parts of capillary loops within the glomerulus are covered by podocytes without fluorescent labeling (Fig. 5a-a”). The same was true for AKS prepared from iPod^Lifeact.mScarlet-I^ animals (Fig. 5b-b”). STED imaging of iPod^Lifeact.mScarlet-I^ shows labeling of single podocytes. The high contrast allows for visualization of an individual podocyte with its foot processes (Fig. 5 c”). Control iPod^Lifeact.mScarlet-I^ animals without tamoxifen induction showed only rarely Lifeact.mScarlet-I positive cells as a sign of a slight leakiness of the inducible promoter (Suppl. Fig. S4).

**Figure 5.**
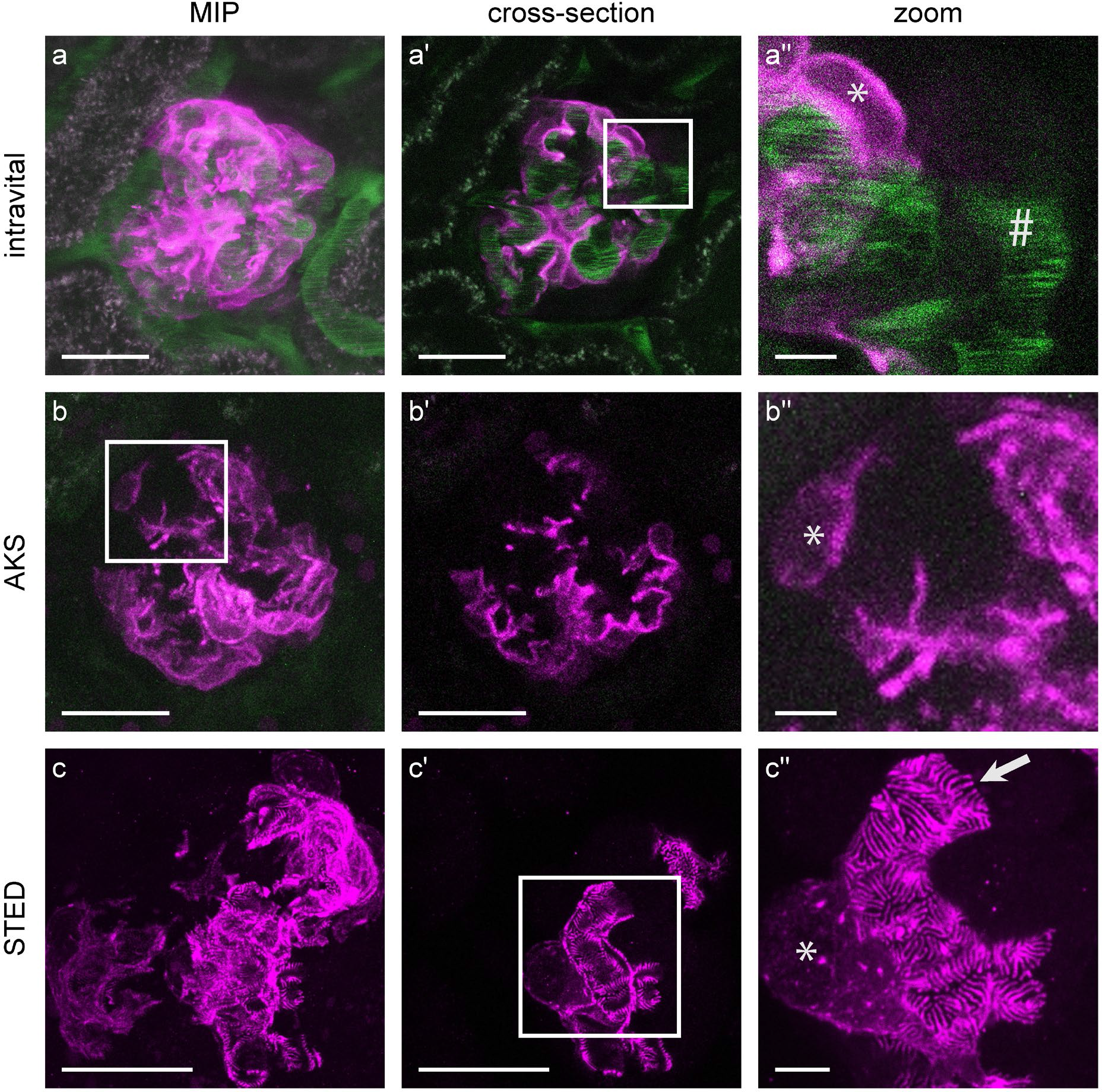
Tamoxifen induced partial expression of Lifeact.mScarlet-I in glomerular podocytes. Representative images of Lifeact.mScarlet-I fluorescence (magenta) in podocytes of an iPod^Lifeact.mScarlet-I^ mouse imaged by intravital microscopy using a multiphoton microscope. Blood flow is visualized by injection of 75-kDa FITC coupled dextran (green). **(b)** Murine AKS of an iPod^Lifeact.mScarlet-I^ animal imaged with multiphoton microscopy. **(c)** Immunofluorescently labeled Lifeact.mScarlet-I tissue section imaged with STED microscopy. All animals received a single dose of tamoxifen via oral gavage (0.1mg/g body weight) and were sacrificed 5 days later. Asterisks indicate podocyte cell bodies. Hashtag # - glomerular capillary; Arrow – single podocyte foot processes; Scale bars – 25 µm, zoom – 5 µm; MIP – maximum intensity projection; AKS – acute kidney slice.

Additionally, we co-stained Lifeact.mScarlet-I with proteins that are important for podocyte ultrastructure, such as acetylated tubulin, vimentin and nephrin. Tubulin is the main cytoskeletal component of primary processes in podocytes and we were interested in the localization of these cytoskeletal proteins with regard to actin fibers. Co-staining of acetylated tubulin and Lifeact.mScarlet-I in iPod^Lifeact.mScarlet-I^ kidney tissue clearly shows that tubulin is exclusively expressed in primary processes, while actin is located in FPs (Fig. 6a-a”). Vimentin, a type of intermediate filament, was mostly present at the base of the cell body and in primary processes, while Lifeact-mScarlet-I staining was mostly limited to the FPs (Fig. 6b+b”). The co-staining with nephrin revealed the localization of the slit diaphragm along the actin fingers in FPs (Fig. 6c-c”). Orthogonal views of a capillary walls cross-section show an alternating nephrin-Lifeact.mScarlet-I-nephrin pattern followed by an empty space of an unlabeled FP (Fig. 6d, XY and XZ view). We then used a transgenic mouse expressing cytosolic GFP in addition to Lifeact.mScarlet-I in podocytes. GFP was evenly distributed in the cytosol, while Lifeact.mScarlet-I was most prominent in FPs (Fig. 6e-e”).

**Figure 6.**
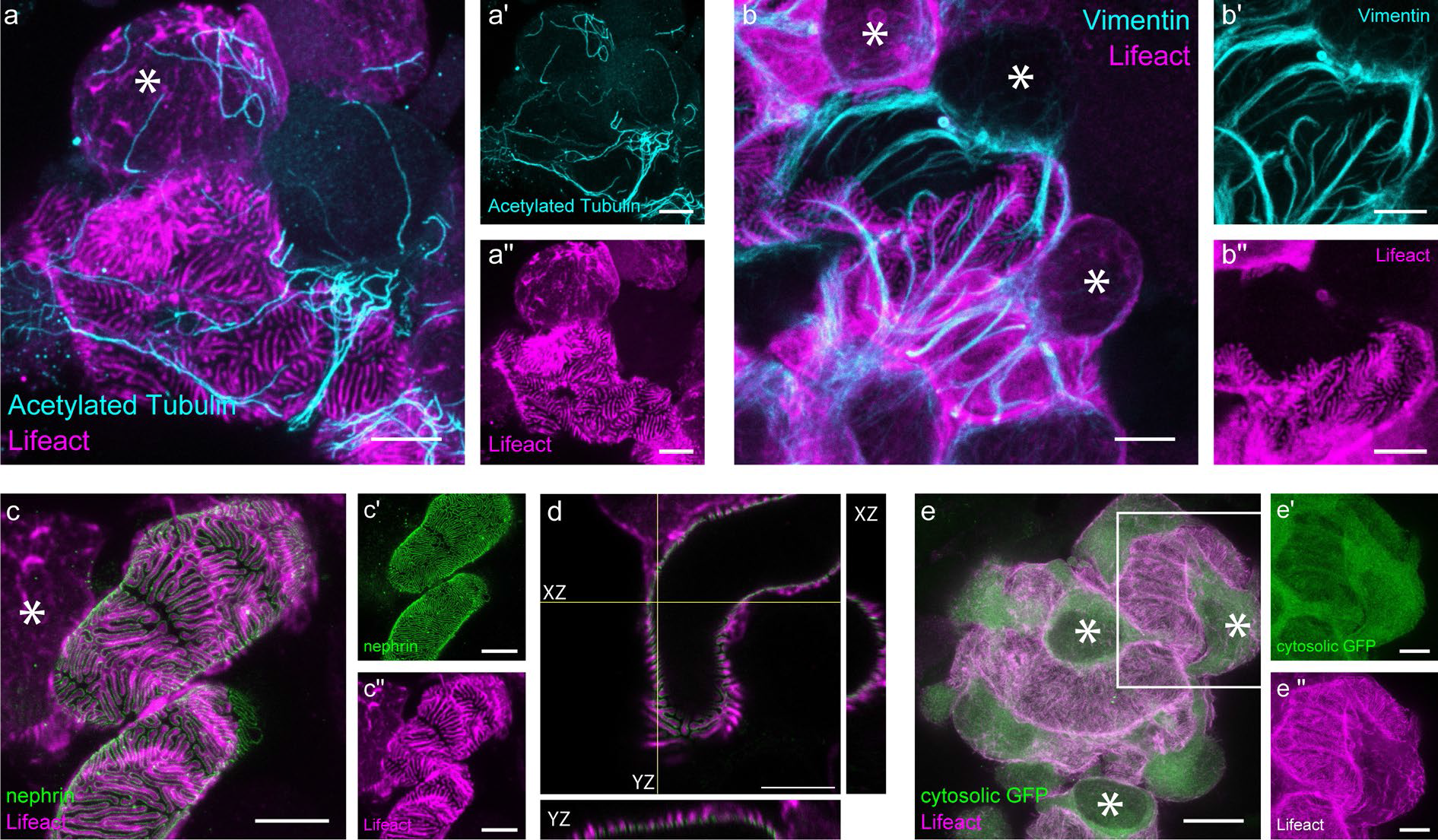
Immunofluorescent stainings of podocyte cytoskeletal components. **(a)** Merged image of a single podocyte of an iPod^Lifeact.mScarlet-I^ mouse stained for acetylated tubulin (cyan) and Lifeact (magenta). Single channels are shown for acetylated tubulin in a’ and Lifeact in a”. Labelling of intermediate filaments by vimentin staining (cyan) and Lifeact (magenta) in a mouse glomerulus of an iPod^Lifeact.mScarlet-I^ animal. Single channels are shown for vimentin in b’ and Lifeact in b”. **(c)** Merged STED image of nephrin (green) and Lifeact (magenta) of an iPod^Lifeact.mScarlet-I^ mouse. Single colors are shown for neprhin in c’ and Lifeact in c”. **(d)** Orthogonal views of foot processes covered glomerular capillary surface with Lifeact (magenta) and nephrin (green) of an iPod^Lifeact.mScarlet-I^ mouse. **(e)** Staining of a cytosolic GFP (green) and Lifeact (magenta) in a Pod^Lifeact.mScarlet-I^ mouse glomerulus. Single colors are shown for GFP in e’ and Lifeact in e”. All iPod^Lifeact.mScarlet-I^ animals received a single dose of tamoxifen via oral gavage (0.1mg/g body weight) and were sacrificed 5 days later. All images are maximum intensity projections except **d**. Asterisks mark podocyte cell bodies. All scale bars – 5 µm, except **e** – 10 µm.

Taken together, this new Lifeact.mScarlet-I mouse line represents a powerful tool to visualize the actin cytoskeleton in podocytes *in vivo, ex vivo* and in STED microscopy in a resolution previously only obtainable by electron microscopy.

## Discussion

Here we present a novel mouse model for tissue-specific expression of Lifeact.mScarlet-I. The reporter is bright, photo stable and suitable for intravital and *ex vivo* imaging to monitor actin structures. Expression was limited to the Cre expressing cells since we did not observe mScarlet-I positive cells within other kidney structures such as tubular cells or endothelial cells. It is therefore plausible that expression of Lifeact.mScarlet-I is also specific using other Cre drivers. Thus, our line is a promising universal tool to visualize F-actin in any tissue desired.

TLA sequencing revealed the correct insertion of a single copy of the Lifeact.mScarlet-I construct into the Gt(ROSA)26Sor locus with no mutations, inversions, or unspecific integration. Phenotypisation of young Pod^Lifeact.mScarlet-I^ and Pod^wt^ animals for general kidney function showed no proteinuria, but higher serum creatinine levels than in older animals. This increase observed in both Pod^wt^ and Pod^Lifeact.mScarlet-I^ genotypes seems to be independent of Cre activity and could be due to diet biases and measurement errors.[25] Analysis of SD length/area morphology in young 4-week-old animals showed a small but significant reduction between Pod^Lifeact.mScarlet-I^ compared to Pod^wt^. This decrease is not typically associated with any changes in proteinuria or histology. Older Pod^Lifeact.mScarlet-I^ animals developed mild proteinuria that correlates with a further reduction of the SD length. This may be caused by interference of Lifeact binding with physiological actin turnover. Intravital imaging experiments are only performed with young animals, and our data show that heterozygous Lifeact.mScarlet-I expression is not affecting kidney function at this age. Due to the bright fluorescence of mScarlet-I, homozygous expression is not necessary for sufficient fluorescence intensity. The effect of Pod^Lifeact.mScarlet-I^ expression on the kidney filter became even more pronounced with age. In line with our results, recently published mouse models such as the beta-actin-GFP mouse line also showed the development of early onset proteinuria and glomerulosclerosis upon aging.[26]

In Pod^Lifeact.mScarlet-I^ animals we observed a homogeneous labeling of the podocyte actin cytoskeleton which is mainly localized in the foot processes and barely in the cell body. The expression pattern of mScarlet-I co-localized with synaptopodin, an actin-binding protein, which is commonly used to indirectly label actin in podocytes in fixed tissue.[27] However, labeling of actin fibers or synaptopodin in all podocytes impaired visualization of single FPs. We therefore used a tamoxifen inducible mouse model to overcome this issue. In iPod^Lifeact.mScarlet-I^ animals the application of a single low dose tamoxifen achieved visualization of single mScarlet-I positive podocytes 5 days later. In these podocytes, the fine and complex structure of FPs could be visualized in detail. Control animals that received only sesame oil did show a low number of mScarlet-I positive cells, indicating minor leakiness of the Pod-ERT2-Cre line.

To demonstrate that our mouse model is capable of visualizing actin changes, we incubated AKS with the cholesterol-depleting agent MβCD. Before treatment, actin was mainly located within the FPs and showed an even distribution. After incubation, rearrangements of the actin cytoskeleton were visible using multiphoton microscopy. Actin was localized more in the cell body and showed actin blebbing.[28] The effect of MβCD was even more striking in super-resolution microscopy. The fine FPs were no longer covering the capillary completely. The subsequent loss of podocytes destroyed the SD partially.

Combining different disease models such as nephrotoxic serum nephritis or adriamycin nephropathy with the Pod^Lifeact.mScarlet-I^ and iPod^Lifeact.mScarlet-I^ mouse model will allow us to study the effects on the podocyte actin cytoskeleton *in vivo* and *ex vivo* upon podocyte injury. The gold standard to visualize podocyte cytoskeletal structures has been electron microscopy.[29] However, the serial sectioning techniques are extremely time consuming and do not allow for co-staining of additional target proteins to give a comprehensive understanding of changes in FP morphology. Our mouse model will now allow putting pathological changes in nephrin patterns into the context of the reorganization of actin and other components of the cytoskeleton such as tubulin and vimentin.

Furthermore, our new mouse models can be combined with genetic disease models including mutations of the SD proteins to understand the structural rearrangements resulting in podocyte FP effacement and focal segmental glomerulosclerosis. Finally, due to its red fluorescence, Lifeact.mScarlet-I is compatible with commonly used green fluorescent reporters such as GCaMPs. The combination of both reporters would allow us to understand the interplay between calcium signaling and actin *in vivo*.

In conclusion, due to its monomeric nature and high fluorescence Lifeact.mScarlet-I is the ideal reporter protein to label actin structures with minimal side effects in podocytes *in vivo, ex vivo* and in fixed tissue. It will allow us to understand how actin rearranges during disease, especially during FP effacement.

## Data availability

The datasets generated, including images acquired with multiphoton or STED microscopy, and analyzed during the current study are available in the figshare repository with the reserved DOI: 10.6084/m9.figshare.26797678.

## Ethical statement

All animal protocols were performed in compliance with the European, national and institutional guidelines and approved by the State Office of North Rhine-Westphalia, Department of Nature, Environment and Consumer Protection (LANUV NRW, Germany; animal study protocol AZ 81-02.04.2019.A335, AZ 81-02.04.2018.A351, AZ 81-02.04.2023.A301).

Acknowledgments

We thank S. Keller, S. Greco-Torres and A. Köser for technical assistance. The authors acknowledge the support of the CECAD Imaging Facility and *in vivo* Research Facility and their Transgenic Core Unit for excellent mouse care and the generation of transgenic mice.

## Author contributions

Multiphoton and STED imaging experiments were designed by J.B.-L. and M.J.H.. S.E.T. and B.Z. generated the mouse model. Experiments were performed by E.W., J.B-L.. The manuscript was written by E.W., J.B-L, N.R. and M.J.H.. T.B., R.W-S., D.U-J. and M.J.H. provided guidance and resources.

## Reference

[1] K. Ichimura, H. Kurihara, and T. Sakai, “Actin filament organization of foot processes in vertebrate glomerular podocytes,” Cell Tissue Res, vol. 329, no. 3, pp. 541–557, 2007, doi: 10.1007/s00441-007-0440-4.

[2] J. K. J. Deegens et al., “Podocyte foot process effacement as a diagnostic tool in focal segmental glomerulosclerosis,” Kidney Int, vol. 74, no. 12, pp. 1568–1576, 2008, doi: 10.1038/ki.2008.413.

[3] J. M. Kaplan et al., “Mutations in ACTN4, encoding α-actinin-4, cause familial focal segmental glomerulosclerosis,” Nat Genet, vol. 24, no. 3, pp. 251–256, 2000, doi: 10.1038/73456.

[4] G. Mollet et al., “Podocin Inactivation in Mature Kidneys Causes Focal Segmental Glomerulosclerosis and Nephrotic Syndrome,” Journal of the American Society of Nephrology, vol. 20, no. 10, pp. 2181–2189, Oct. 2009, doi: 10.1681/ASN.2009040379.

[5] J. H. Kim et al., “CD2-associated protein haploinsufficiency is linked to glomerular disease susceptibility,” Science (1979), vol. 300, no. 5623, pp. 1298–1300, May 2003, doi: 10.1126/science.1081068.

[6] J. Riedl et al., “Lifeact: A versatile marker to visualize F-actin,” Nat Methods, vol. 5, no. 7, pp. 605–607, 2008, doi: 10.1038/nmeth.1220.

[7] J. Riedl et al., “Lifeact mice for studying F-actin dynamics,” Nat Methods, vol. 7, no. 3, pp. 168–169, 2010, doi: 10.1038/nmeth0310-168.

[8] H. Schachtner et al., “Tissue inducible Lifeact expression allows visualization of actin dynamics in vivo and ex vivo,” Eur J Cell Biol, vol. 91, no. 11–12, pp. 923–929, 2012, doi: 10.1016/j.ejcb.2012.04.002.

[9] K. Sliogeryte, S. D. Thorpe, Z. Wang, C. L. Thompson, N. Gavara, and M. M. Knight, “Differential effects of LifeAct-GFP and actin-GFP on cell mechanics assessed using micropipette aspiration,” J Biomech, vol. 49, no. 2, pp. 310–317, Jan. 2016, doi: 10.1016/j.jbiomech.2015.12.034.

[10] A. Kumari, S. Kesarwani, M. G. Javoor, K. R. Vinothkumar, and M. Sirajuddin, “Structural insights into actin filament recognition bycommonly used cellular actin markers,” EMBO J, vol. 39, no. 14, p. e104006, 2020, doi: 10.1101/846337.

[11] H. Mazloom-Farsibaf, F. Farzam, M. Fazel, M. J. Wester, M. B. M. Meddens, and K. A. Lidke, “Comparing Lifeact and Phalloidin for Super-resolution Imaging of Actin in Fixed Cells,” PLoS One, vol. 16, no. 1, p. e.0246138, 2021, doi: 10.1371/journal.pone.0246138.

[12] W. Wegner et al., “In vivo mouse and live cell STED microscopy of neuronal actin plasticity using far-red emitting fluorescent proteins,” Sci Rep, vol. 7, no. 1, p. 11781, 2017, doi: 10.1038/s41598-017-11827-4.

[13] D. S. Bindels et al., “MScarlet: A bright monomeric red fluorescent protein for cellular imaging,” Nat Methods, vol. 14, no. 1, pp. 53–56, 2016, doi: 10.1038/nmeth.4074.

[14] D. Unnersjö-Jess, L. Scott, H. Blom, and H. Brismar, “Super-resolution stimulated emission depletion imaging of slit diaphragm proteins in optically cleared kidney tissue,” Kidney Int, vol. 89, no. 1, pp. 243–247, 2016, doi: 10.1038/ki.2015.308.

[15] N. Artelt et al., “Comparative analysis of podocyte foot process morphology in three species by 3D super-resolution microscopy,” Front Med (Lausanne), vol. 5, no. 292, p. eCollection 2018, 2018, doi: 10.3389/fmed.2018.00292.

[16] V. T. Chu et al., “Efficient generation of Rosa26 knock-in mice using CRISPR/Cas9 in C57BL/6 zygotes,” BMC Biotechnol, vol. 16, no. 1, pp. 1–15, 2016, doi: 10.1186/s12896-016-0234-4.

[17] S. E. Tröder, L. K. Ebert, L. Butt, S. Assenmacher, B. Schermer, and B. Zevnik, “An optimized electroporation approach for efficient CRISPR/Cas9 genome editing in murine zygotes,” PLoS One, vol. 13, no. 5, p. e.0196891, 2018, doi: 10.1371/journal.pone.0196891.

[18] M. Wigger, M. Schneider, A. Feldmann, S. Assenmacher, B. Zevnik, and S. E. Tröder, “Successful use of HTF as a basal fertilization medium during SEcuRe mouse in vitro fertilization,” BMC Res Notes, vol. 16, no. 184, 2023, doi: 10.1186/s13104-023-06452-6.

[19] J. Wang et al., “Tamoxifen-inducible podocyte-specific iCre recombinase transgenic mouse provides a simple approach for modulation of podocytes in vivo,” Genesis, vol. 48, no. 7, pp. 446–451, 2010, doi: 10.1002/dvg.20635.

[20] M. J. Moeller, S. K. Sanden, A. Soofi, R. C. Wiggins, and L. B. Holzman, “Podocyte-specific expression of Cre recombinase in transgenic mice,” Genesis (United States), vol. 35, no. 1, pp. 39–42, Jan. 2003, doi: 10.1002/gene.10164.

[21] G. E. Truett, P. Heeger, R. L. Mynatt, A. A. Truett, J. A. Walker, and M. L. Warman, “Preparation of PCR-quality mouse genomic dna with hot sodium hydroxide and tris (HotSHOT),” Biotechniques, vol. 29, no. 1, pp. 52–54, Jul. 2000, doi: 10.1016/j.jacc.2014.05.073.

[22] J. Binz-lotter et al., “Injured Podocytes Are Sensitized to Angiotensin II – Induced Calcium Signaling,” Journal of the American Journal of Nephrology, vol. 31, no. 3, pp. 532–542, 2020, doi: 10.1681/ASN.2019020109.

[23] E. Wiesner et al., “Correlative multiphoton-STED microscopy of podocyte calcium levels and slit diaphragm ultrastructure in the renal glomerulus,” Sci Rep, vol. 14, no. 1, Dec. 2024, doi: 10.1038/s41598-024-63507-9.

[24] F. Poosti et al., “Precision-cut kidney slices (PCKS) to study development of renal fibrosis and efficacy of drug targeting ex vivo,” The Company of Biologists, vol. 8, pp. 1227–1236, 2015.

[25] P. Meneton, I. Ichikawa, T. Inagami, and J. Schnermann, “Renal physiology of the mouse,” American Journal of Physiology-Renal Physiology, vol. 278, no. 3, pp. F339–F351, Mar. 2000, doi: 10.1152/ajprenal.2000.278.3.F339.

[26] J. K. Guo et al., “The commonly used β-actin-GFP transgenic mouse strain develops a distinct type of glomerulosclerosis,” Transgenic Res, vol. 16, no. 6, pp. 829–834, Dec. 2007, doi: 10.1007/s11248-007-9107-x.

[27] H. Y. Suleiman, R. Roth, S. Jain, J. E. Heuser, A. S. Shaw, and J. H. Miner, “Injury-induced actin cytoskeleton reorganization in podocytes revealed by super-resolution microscopy,” JCI Insight, vol. 2, no. 16, Aug. 2017, doi: 10.1172/JCI.INSIGHT.94137.

[28] G. R. Wickman et al., “Blebs produced by actin-myosin contraction during apoptosis release damage-associated molecular pattern proteins before secondary necrosis occurs,” Cell Death Differ, vol. 20, no. 10, pp. 1293–1305, Oct. 2013, doi: 10.1038/cdd.2013.69.

[29] K. Ichimura et al., “Morphological process of podocyte development revealed by block-face scanning electron microscopy,” J Cell Sci, vol. 130, no. 1, pp. 132–142, 2017, doi: 10.1242/jcs.187815.

