## Supplementary material for "A Red Fluorescent Lifeact Marker to Study Actin Morphology in Podocytes"

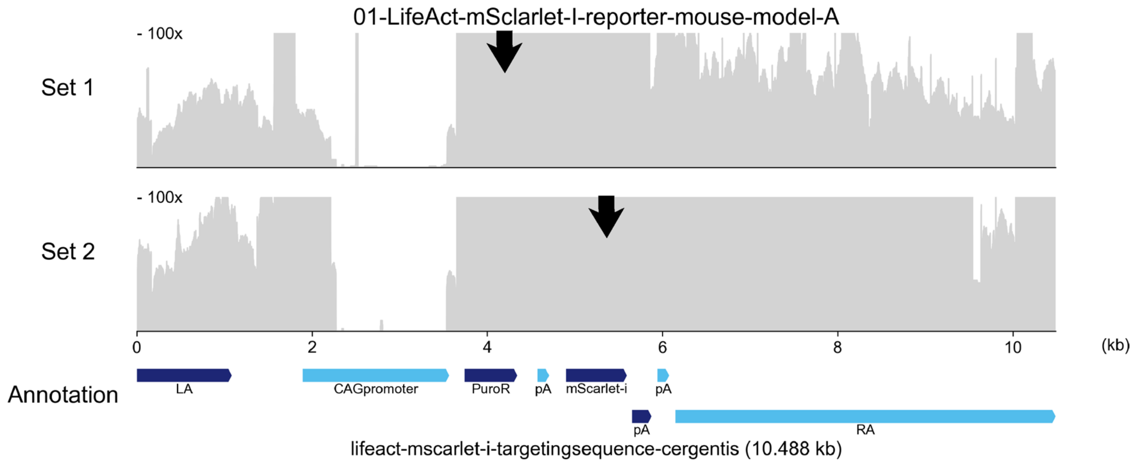


**Supplementary Figure S1. Targeted locus amplification sequencing for validation of correct Lifeact.mScarlet-I integration into the R26 locus.** NGS sequencing coverage (in grey) across the Lifeact.mScarlet-I vector. Black arrows indicate the primer binding location. The annotated vector map is shown at the bottom. Y-axes are limited to 100x. Annotation: LA – Left homology arm, PuroR – Puromycin Resistance, pA – Poly A tail, RA – Right homology arm.

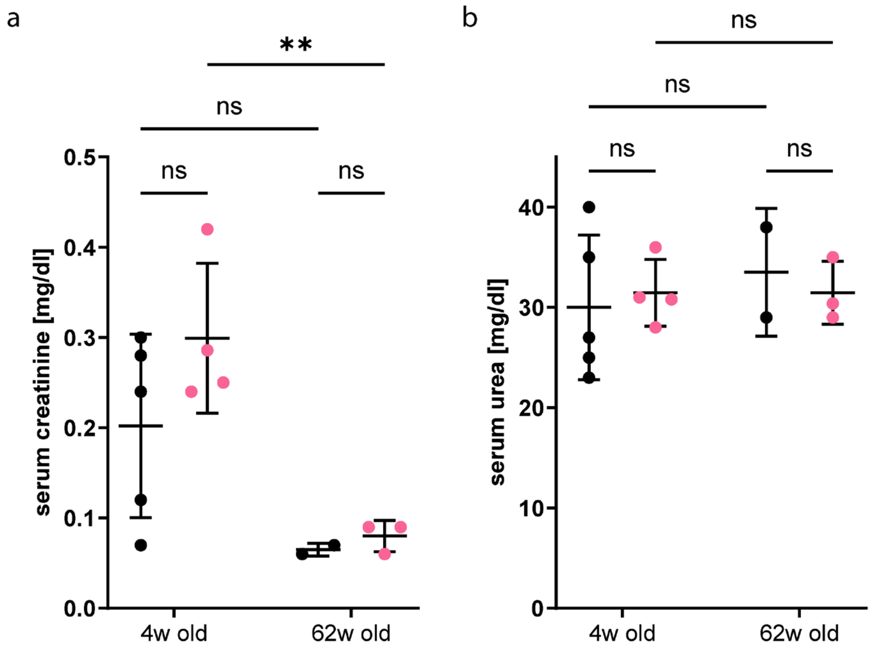


**Supplementary Figure S2. Serum urea and creatinine levels measured in young and aged Pod$^{\text{Lifeact.mScarlet-I}}$. (a)** Serum creatinine levels of young and aged Pod$^{\text{Lifeact.mScarlet-I}}$ mice. **(b)** Serum urea levels of young and aged Pod$^{\text{Lifeact.mScarlet-I}}$ mice. In both measurements the following animals were analyzed: Pod$^{\text{Lifeact.mScarlet-I}}$ 4-weeks: n=4; 62 weeks: n=3 and Pod$^{\text{wt}}$: 4-weeks: n=5; 62 weeks n=2. Statistical analysis: Ordinary two-way ANOVA with multiple comparisons. ns $p > 0.05$, * $p \leq 0.05$, ** $p \leq 0.01$, *** $p \leq 0.001$, **** $p \leq 0.0001$.

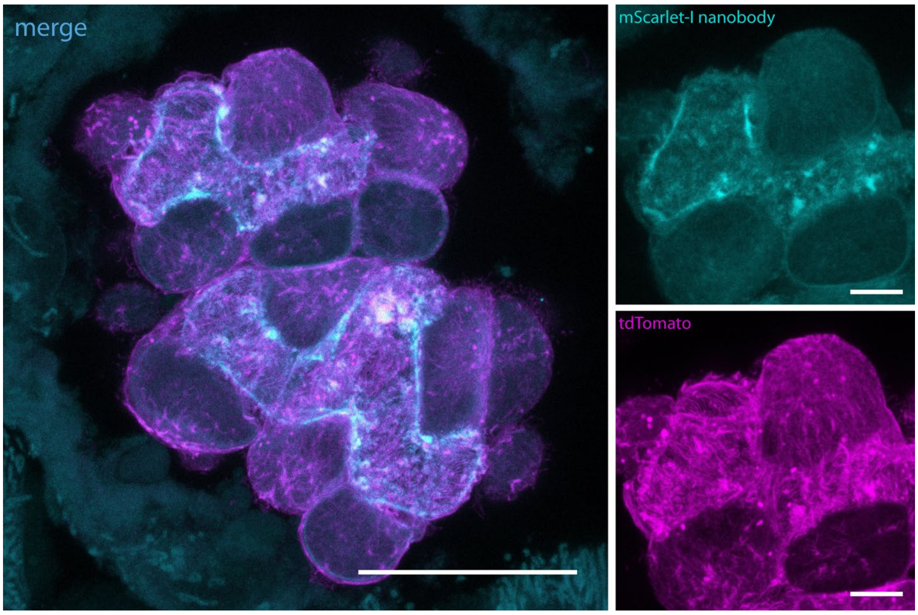


**Supplementary Figure S3. Staining of m.Scarlet-I using an anti-mScarlet-I nanobody.** MIP image of a single glomerulus of Pod$^{Lifeact.mScarlet-I}$ mouse tissue stained for Lifeact.mScarlet-I using an anti mScarlet-I nanobody (cyan) and an anti tdTomato antibody (magenta). Overlap appears in blue. Scale bars – 10 µm, zoom – 5µm.

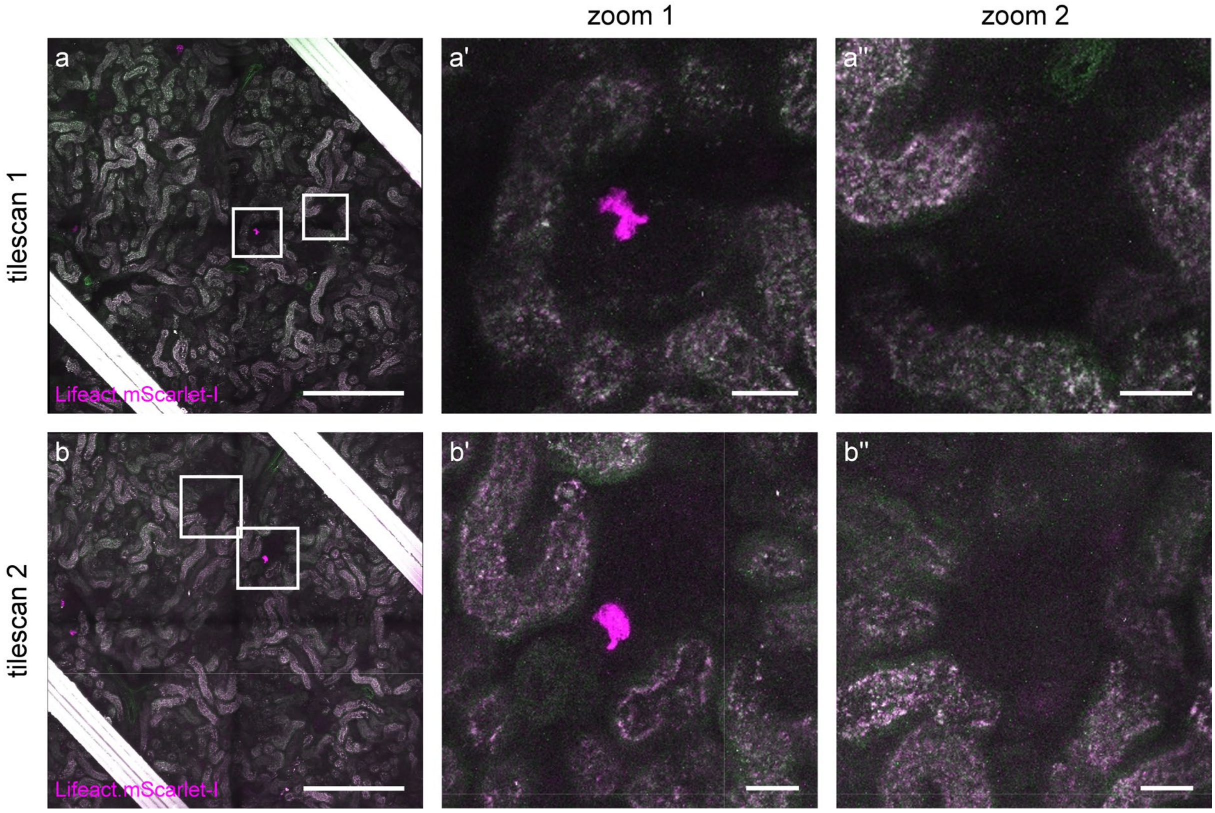


**Supplementary Figure S4. Control iPod$^{Lifeact.mScarlet-I}$ mice induced with sesame oil show low number of mScarlet-I positive cells**. Representative multiphoton tilescan images of AKS of iPod$^{Lifeact.mScarlet-I}$ mouse induced with only sesame oil and no tamoxifen and imaged 5 days later using multiphoton microscopy. **(a)**, **(b)** Tilescan 1 and 2 give an overview of a larger tissue section. Zoom ins (a',a'' and b', b'') highlight glomeruli with (zoom 1) and without (zoom 2) mScarlet-I positive cells. Scale bars – 300 µm, zoom – 25 µm.

**Supplementary Video V1. Intravital microscopy of iPod$^{Lifeact.mScarlet-I}$ mouse glomerulus using multiphoton microscopy.** Z-Stack video of glomerulus with induced expression of Lifeact.mScarlet-I (magenta) in podocytes. Blood flow was labeled by i.a. injection of FITC-Dextran. Scale bar – 25 µm

**Supplementary Video V2. 3D projection of iPod$^{Lifeact.mScarlet-I}$ mouse glomerulus using STED microscopy.** Z-stack shows nephrin staining (green) and Lifeact.mScarlet-I (magenta) of a single podocyte covering a glomerular capillary. The primary process lacks Lifeact.mScarlet-I signal.
